# Mushroom body input connections form independently of sensory activity in *Drosophila melanogaster*

**DOI:** 10.1101/2021.11.06.467552

**Authors:** Tatsuya Hayashi, Alexander John MacKenzie, Ishani Ganguly, Hayley Marie Smihula, Miles Solomon Jacob, Ashok Litwin-Kumar, Sophie Jeanne Cécile Caron

## Abstract

Associative brain centers, such as the insect mushroom body, need to represent sensory information in an efficient manner. In *Drosophila melanogaster*, the Kenyon cells of the mushroom body integrate inputs from a random set of olfactory projection neurons, but some projection neurons — namely those activated by a few ethologically meaningful odors — connect to Kenyon cells more frequently than others. This biased and random connectivity pattern is conceivably advantageous, as it enables the mushroom body to represent a large number of odors as unique activity patterns while prioritizing the representation of a few specific odors. How this connectivity pattern is established remains largely unknown. Here, we test whether the mechanisms patterning the connections between Kenyon cells and projection neurons depend on sensory activity or whether they are hardwired. We mapped a large number of mushroom body input connections in anosmic flies — flies lacking the obligate odorant co-receptor Orco — and in wildtype flies. Statistical analyses of these datasets reveal that the random and biased connectivity pattern observed between Kenyon cells and projection neurons forms normally in the absence of most olfactory sensory activity. This finding supports the idea that even comparatively subtle, population-level patterns of neuronal connectivity can be encoded by fixed genetic programs and are likely to be the result of evolved prioritization of ecologically and ethologically salient stimuli.

## INTRODUCTION

The precise wiring between sensory systems and higher brain centers is orchestrated by a combination of hardwired and activity-dependent mechanisms [1]. Hardwired mechanisms include complex signaling networks that guide neuronal outgrowths to their target and cell surface molecules that pair synaptic partners. Such hardwired mechanisms are necessary to establish coarse connectivity patterns in a reproducible and reliable manner. In contrast, activity-dependent mechanisms — sensory or spontaneous activity — can refine these coarse patterns by promoting the connectivity of active inputs over that of inactive inputs. While spontaneous activity can refine connectivity patterns independently of experience, sensory activity sculpts connections based on available information such that the overall structure of a network can be molded after the sensory environment peculiar to an organism.

The *Drosophila melanogaster* mushroom body is a higher brain center formed by 2,000 neurons called ‘Kenyon cells’ and primarily processes olfactory information [2,3]. The primary olfactory center in the fly brain, the antennal lobe, consists of 51 glomeruli; each glomerulus receives input from a set of olfactory sensory neurons expressing the same receptor gene(s) [4,5]. Olfactory information is relayed from individual glomeruli to Kenyon cells by about 160 uniglomerular projection neurons [6,7]. The connections between projection neurons and Kenyon cells are random: individual Kenyon cells integrate inputs from a small set of projection neurons that cannot be assigned to a common group based on their biological characteristics [3,8–10]. Such a random connectivity pattern has been predicted by several theoretical studies to be advantageous, as it expands the capacity of the mushroom body to represent olfactory information by minimizing the overlap between representations [11,12].

Although random, the connections between projection neurons and Kenyon cells are also biased: not all projection neurons connect to Kenyon cells at the same frequency — some neurons are overrepresented while others are underrepresented [3,9]. Interestingly, the most biased projection neurons, both underrepresented and overrepresented neurons, receive input from olfactory sensory neurons narrowly tuned to detect odors that are particularly meaningful. For instance, the DP1m and DA1 projection neurons are among the most overrepresented neurons. The DP1m projection neuron receives input from the IR64a-expressing olfactory sensory neurons, which detect acids produced by fermenting fruits, a potential food source, whereas the DA1 projection neurons receive input from the OR67d expressing neurons which detect the pheromone 11-*cis*-vaccenyl acetate [13,14]. In contrast, underrepresented projection neurons are activated by odors that trigger strong innate avoidance, likely via mushroom-body-independent pathways. For instance, the DL4 and DA2 projection neurons are among the most underrepresented neurons. The DL4 projection neuron receives input from the OR49a/OR85f-expressing neurons that detect odors produced by parasitoid wasps whereas the DA2 projection neurons receive input from the OR56a-expressing neurons that detect odors produced by toxic microbes [15,16].

Whether biases in connectivity arise through sensory activity or hardwired mechanisms is not known. To distinguish between these two possibilities, we sought to compare whether the mushroom body input connections differ between wildtype flies and flies in which most olfactory sensory neurons are silent.

## RESULTS

### Decreased odor-evoked activity in the mushroom body calyx of *Orco*^-/-^ flies

Orco — also known as OR83b — is the obligate co-receptor of all Odorant Receptors (ORs) in most insects [17,18]. Orco is required for olfactory transduction, and, hence, OR-expressing neurons in *Orco*^-/-^ flies do not show odor-evoked responses [17]. Of the 51 antennal lobe glomeruli, at least 37 receive input from OR-expressing sensory neurons (Table S1). The remaining glomeruli are innervated by olfactory sensory neurons expressing either Ionotropic Receptors (IRs), which are tuned to amines and acids, or Gustatory Receptors (GRs), which detect carbon dioxide [14,18–20]. Both IRs and GRs do not require Orco as a co-receptor, therefore *Orco*^-/-^ flies are not completely anosmic and can detect odors that bind to these receptors [14]. To test whether these sensory defects are reflected in the mushroom body, we measured odor-evoked responses in the calyx — the neuropil where projection neurons connect with Kenyon cells — of *Orco*^+/+^ and *Orco*^-/-^ flies that express GCaMP6f in all Kenyon cells. As expected, we detected no odor-evoked calcium transients in the calyx of *Orco*^-/-^ flies in response to isopentyl acetate, an odor detected by several ORs, but we detected odor-evoked responses to acetic acid, an odor detected by IRs (Figure 1A,B). In contrast, the calyx of *Orco*^+/+^ flies show large calcium transients in response to both odors (Figure 1A,B). Similar results were obtained across different flies (Figure 1C).

**Figure 1.**
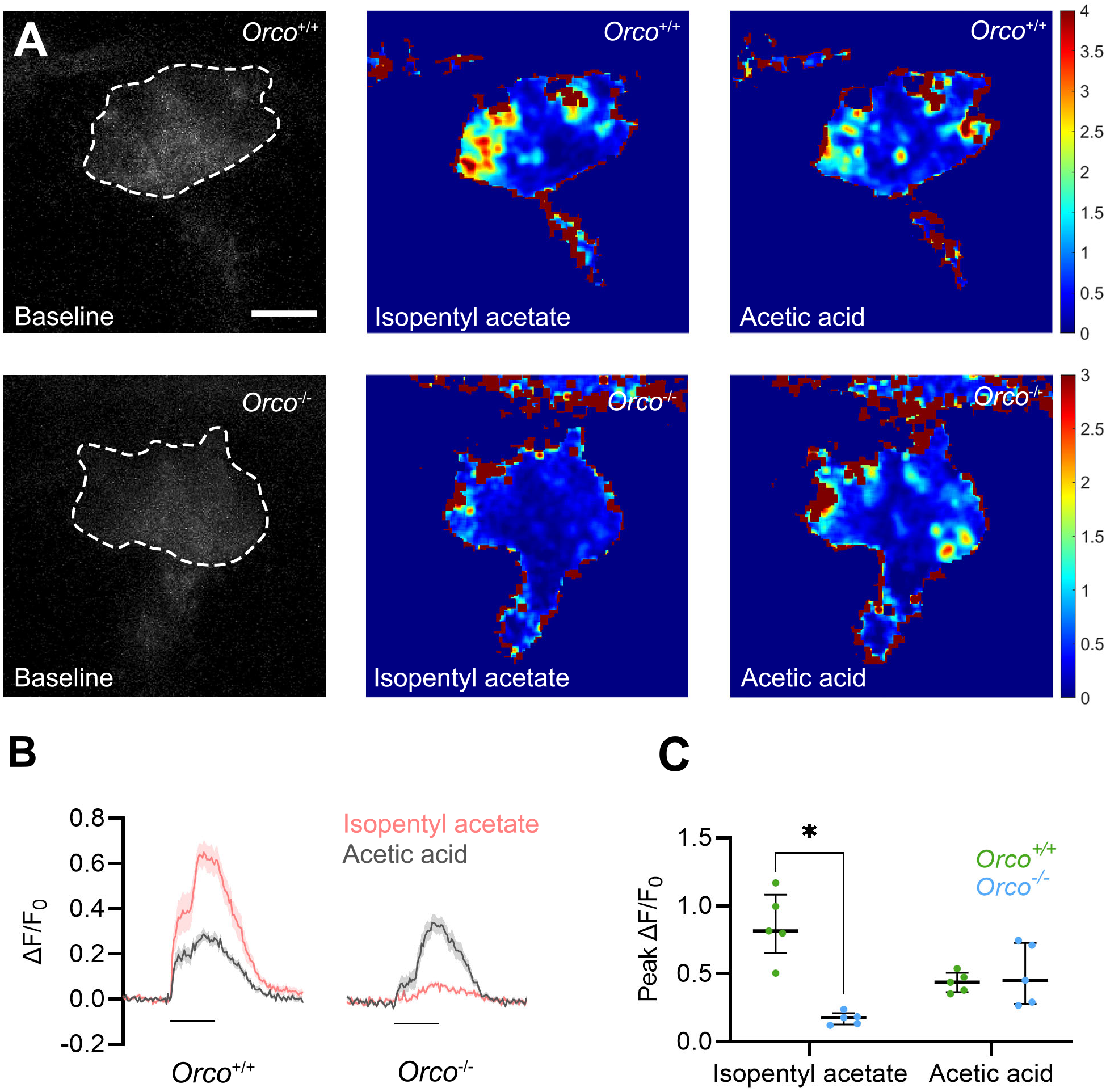
Odor-evoked activity is decreased in the mushroom body calyx of *Orco*^-/-^ flies. (A-C) Calcium imaging in Kenyon cells shows odor-evoked activity in *Orco*^-/-^ flies in response to isopentyl acetate, an odor that activates multiple ORs, but not in response to acetic acid, an odor that activates IRs. (A) Kenyon cells extend their dendrites in the calyx of the mushroom body which can be recognized based on baseline fluorescence in *Orco*^+/+^ and *Orco*^-/-^ flies (left panels, dashed white line). Example heatmaps show ΔF/F_0_ in response to isopentyl acetate (middle panels) and acetic acid (right panels). The color bars denote the range of ΔF/F_0_ in each sample. Scale bar is 20 μm. (B) The ΔF/F_0_ values recorded in the main calyx in response to isopentyl acetate (pink) and acetic acid (gray) were averaged across five trials collected in five different animals and are shown as traces; the shaded area of each trace represents the standard deviation. (C) The peak ΔF/F_0_ values were averaged across trials in each of the five *Orco*^+/+^ (green) and *Orco*^-/-^ (blue) flies; the long black bars represent the median whereas the short black bars represent the 25th and 75th percentile of the data; the asterisk indicates values that were statistically different (*p* < 0.05). The statistical significance, or ‘*p*-value’, was measured using the Mann-Whitney U test.

These results show that sensory activity is severely impaired in the mushroom body of *Orco*^-/-^ flies: while Kenyon cells respond to an odor detected by IR-expressing neurons, they fail to respond to an odor detected by OR-expressing neurons.

### Glomeruli of the antennal lobe are morphologically similar in *Orco*^+/+^ and *Orco*^-/-^ flies

Next, we investigated whether the neuroanatomy of the antennal lobe of *Orco*^-/-^ flies differs from that of *Orco*^+/+^ flies. Previous studies have demonstrated that in *Orco*^-/-^ flies olfactory sensory neurons are able to target their cognate glomeruli [21–23]. However, a recent study showed that in ants *Orco* loss-of-function leads to smaller antennal lobes that contain fewer glomeruli [24,25]. To verify whether similar defects are found in *Orco*^-/-^ flies, we reconstructed their antennal lobes and identified individual glomeruli based on available anatomical maps as well as the hemibrain connectome [4,5,26]. We identified a total of 51 glomeruli in both genotypes (Figure 2A). The volume of individual antennal lobes in *Orco*^+/+^ and *Orco*^-/-^ flies is not significantly different (antennal lobe volume (median): *Orco*^+/+^: 84723 μm^3^, *Orco*^-/-^: 81128 μm^3^, *p*-value = 0.15; Figure 2B and Table S1). Most glomeruli receiving input from OR-expressing neurons are slightly smaller in *Orco*^-/-^ flies whereas some glomeruli receiving input from IR/GR-expressing neurons are slightly larger, but none of these differences are statistically significant (Figure 2C; Table S1).

**Figure 2.**
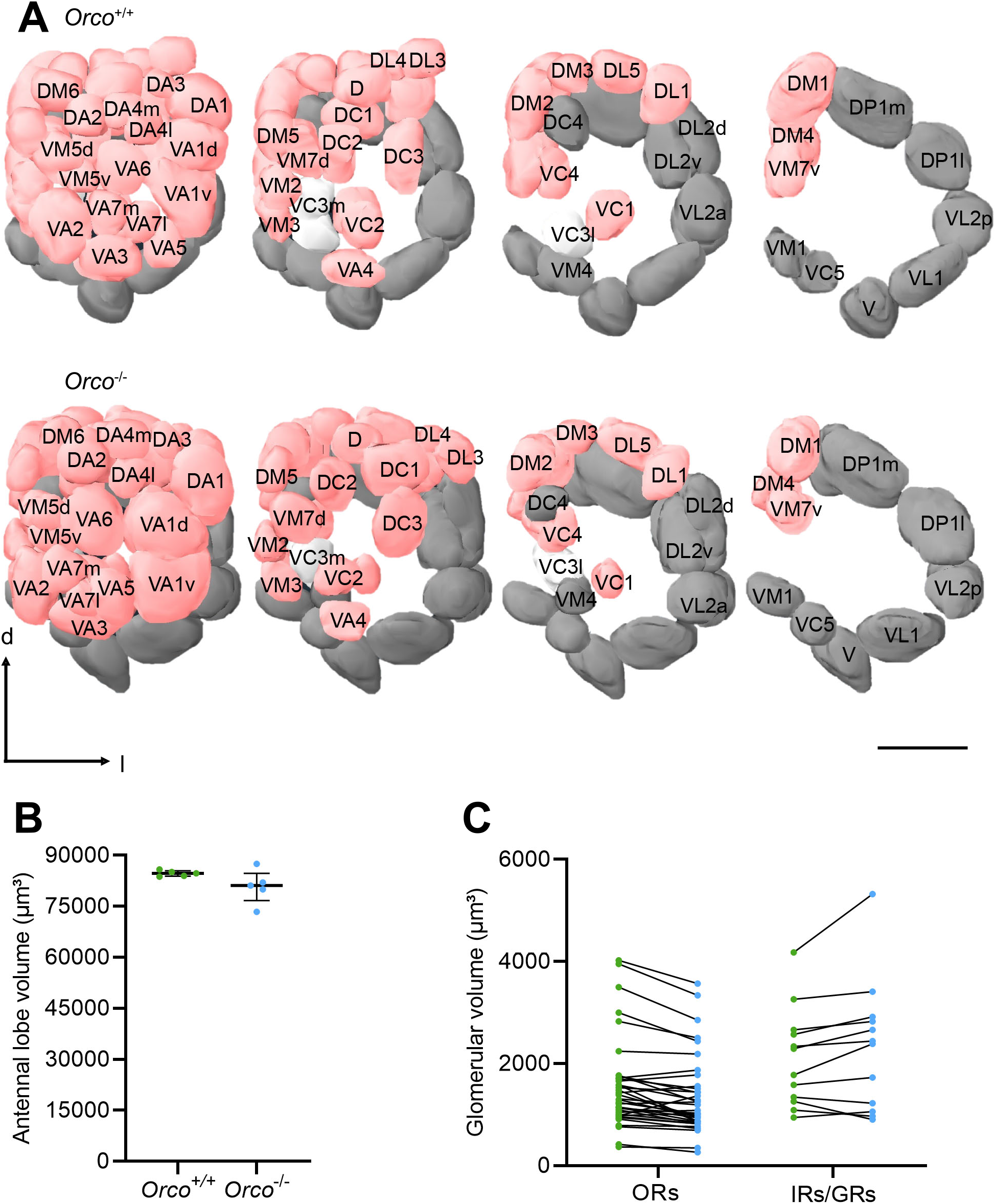
Antennal lobes are morphologically similar in *Orco*^+/+^ and *Orco*^-/-^ flies. (A) The brains of *Orco*^+/+^ and *Orco*^-/-^ flies were fixed, immuno-stained (using anti-bruchpilot also known as nc82) and imaged; each of the 51 glomeruli forming the antennal lobe were reconstructed and identified based on theirs shapes and locations; each glomerulus receives input from either ORs-expressing neurons (pink), IRs/GRs-expressing neurons (dark gray) or unidentified sensory neurons (light gray). Scale bar is 25 μm. (B-C) The reconstructed volumes of the entire antennal lobe (B) or individual glomeruli (C) were compared across genotypes (green: *Orco*^+/+^; blue: *Orco*^-/-^); the long black bars represent the median whereas the short black bars represent the 25th and 75th percentile of the data. (C) The volumes of a given glomerulus in both genotypes are link with a black line, and the statistical significance of the differences in glomerular volumes are provided in Table S1. The statistical significance, or ‘*p*-value’, was measured using the Mann-Whitney U test.

Altogether, these results show that the antennal lobes form normally in the absence of sensory activity: although some glomeruli appear to be slightly smaller or larger in *Orco*^-/-^ flies, all glomeruli were present.

### Projection neurons and Kenyon cells are morphologically similar in *Orco*^+/+^ and *Orco*^-/-^ flies

Next, we investigated whether the projection neurons connecting individual glomeruli to Kenyon cells show morphological differences between *Orco*^+/+^ and *Orco*^-/-^ flies. To this end, we photolabeled the neurons innervating the DL4 glomerulus, which receives input from the OR49a+/OR85f+ neurons, the VA2 glomerulus, which receives input from the OR92a+ neurons, and the DP1m glomerulus, which receives input from the IR64a+ neurons (Figure 3A and Figure S1). For each glomerulus, we recovered a single photo-labeled projection neuron in both genotypes (Figure 3B). We determined the number of presynaptic clusters — or ‘bouton clusters’ — each of these projection neurons form in the mushroom body as well as the number of branches each neuron extends; we also measured the total volume of the boutons formed by a given projection neuron (Figure 3C-E). Based on these measurements, we found that the projection neurons of *Orco*^+/+^ and *Orco*^-/-^ flies are largely comparable. There are some small but significant differences: the DL4 projection neuron forms fewer branches and fewer bouton clusters in *Orco*^-/-^ flies (number of branches (median): *Orco*^+/+^: 2.5, *Orco*^-/-^: 1.5, *p*-value = 0.02; number of bouton clusters (median): *Orco*^+/+^: 3.0, *Orco*^-/-^: 2.0, *p*-value = 0.009; Figure 3C,E) whereas the volume of the boutons formed by the VA2 projection neuron is slightly larger in *Orco*^-/-^ flies (bouton volume (median): *Orco*^+/+^: 422.8 μm^3^, *Orco*^-/-^: 602.3 μm^3^, *p*-value = 0.03; Figure 3E).

**Figure 3.**
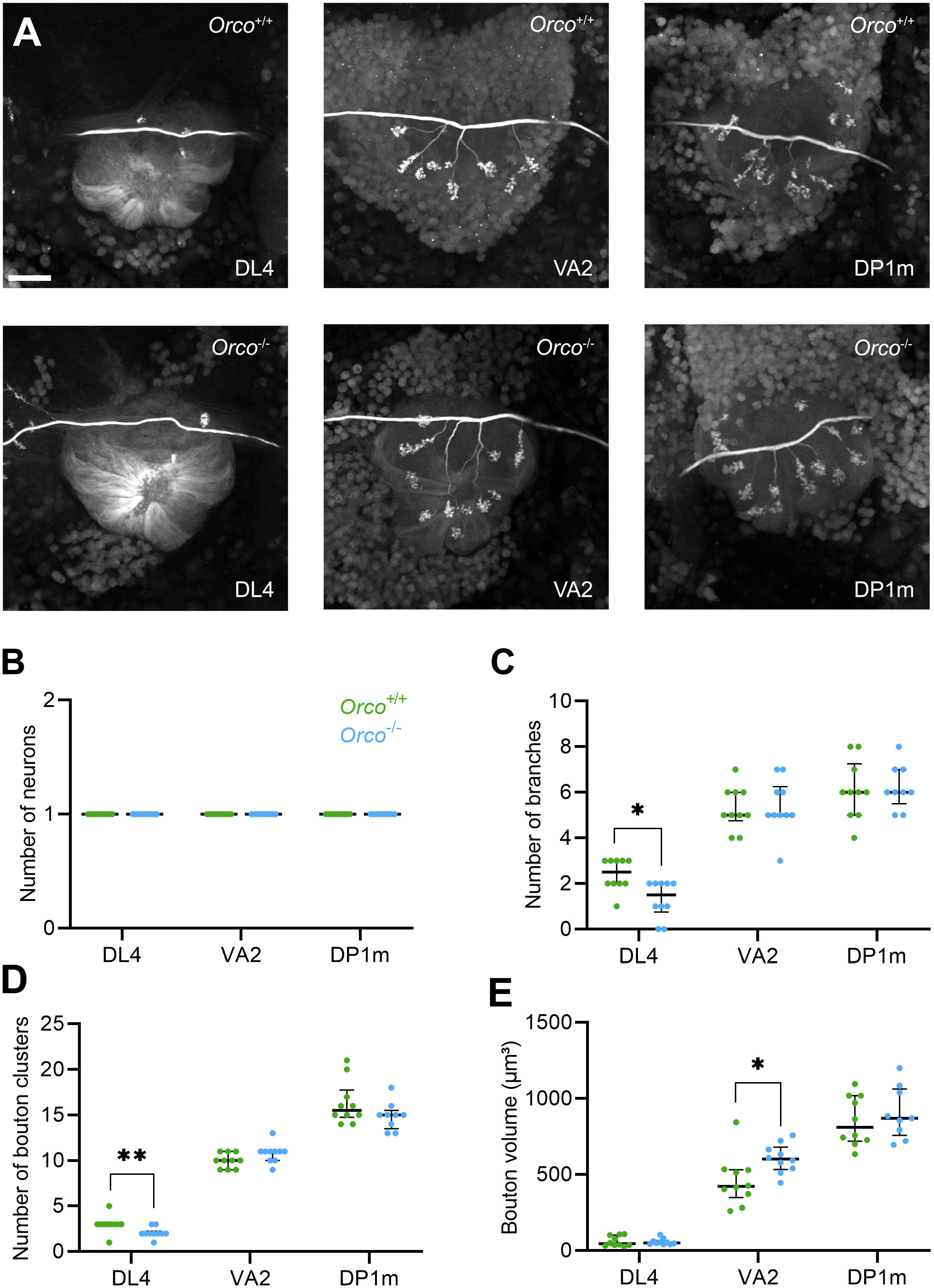
Projection neurons are morphologically similar in *Orco*^+/+^ and *Orco*^-/-^ flies. (A) The projection neurons innervating the DL4 (left), VA2 (middle) and DP1m (right) glomeruli were photo-labeled in *Orco*^+/+^ and *Orco*^-/-^ flies, and the presynaptic terminals — called ‘boutons’ — formed by these neurons in the calyx of the mushroom body were imaged. Scale bar is 15 μm. (B-E) The number of photo-labeled projection neurons was counted (B) as well as the number of bouton clusters (C) and primary branches (D) found in *Orco*^+/+^ (green) and *Orco*^-/-^ flies (blue); the total bouton volume was quantified (E); the long black bars represent the median whereas the short black bars represent the 25th and 75th percentile of the data; the asterisks indicate values that were statistically different (*: *p* < 0.05 and **: *p* < 0.01). The statistical significance, or ‘*p*-value’, was measured using the Mann-Whitney U test.

Next, we determined whether Kenyon cells show morphological differences between *Orco*^+/+^ and *Orco*^-/-^ flies. Based on their axonal projection patterns, Kenyon cells can be divided into three major types: alpha/beta, alpha’/beta’ and gamma Kenyon cells [2]. It has previously been shown that the number of post-synaptic terminals, or ‘claws’, formed by a neuron varies across types [3,9]. We photo-labeled individual Kenyon cells of each type and compared their morphological features between *Orco*^+/+^ and *Orco*^-/-^ genotypes (Figure 4A, Figure S2). Specifically, we measured the total length of the branches individual Kenyon cells extend in the calyx, as well as the number and length of the claws formed by individual Kenyon cells (Figure 4B-D, Figure S2). We detected one significant difference: gamma Kenyon cells form longer branches in *Orco*^-/-^ flies than they do in *Orco*^+/+^ flies (total branch length (median): *Orco*^+/+^: 87.8, *Orco*^-/-^: 147.0, *p*-value = 0.02; Figure 4B). However, apart from this difference, Kenyon cells appear to be morphologically similar in both genotypes.

**Figure 4.**
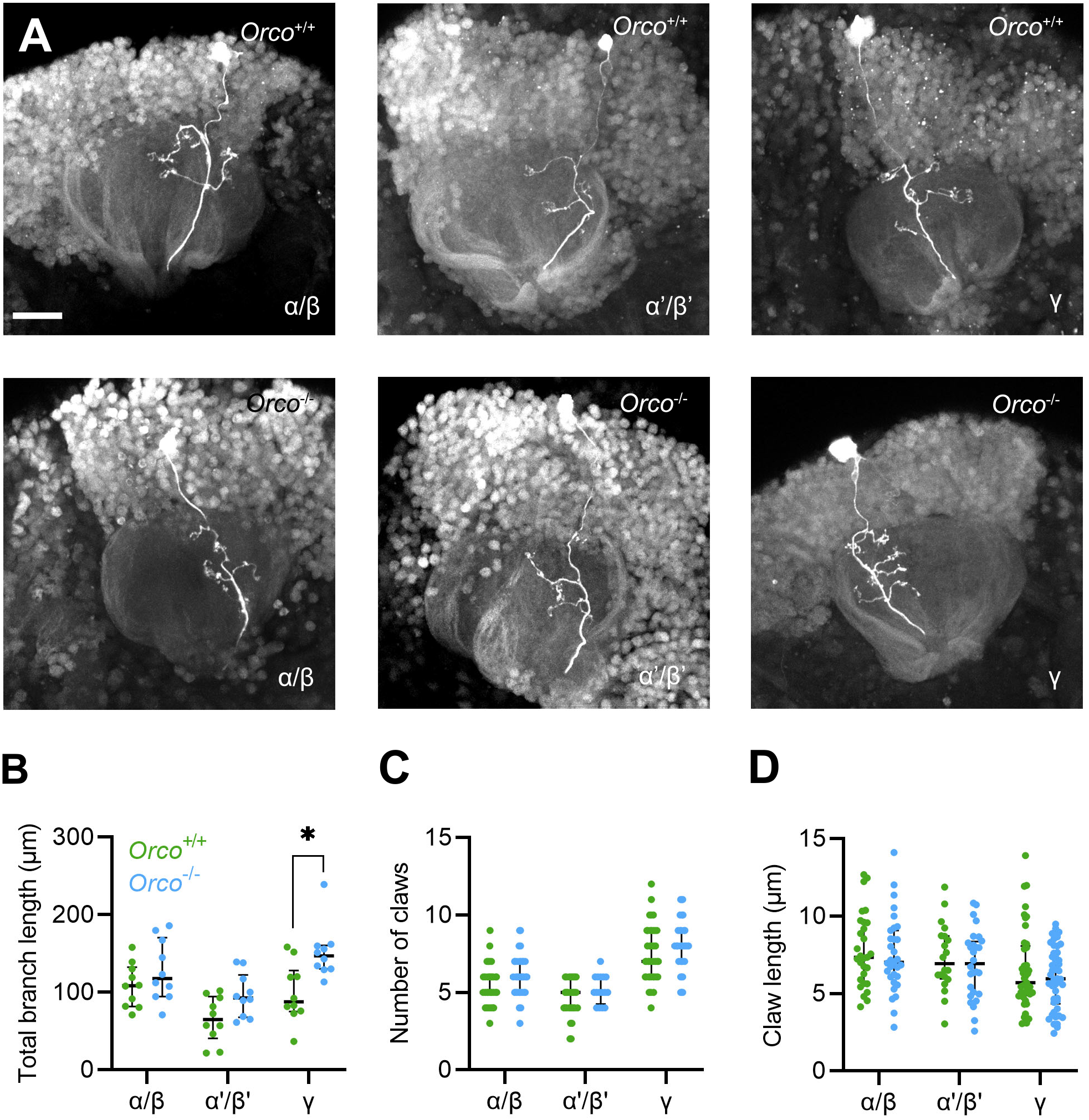
Kenyon cells are morphologically similar in *Orco*^+/+^ and *Orco*^-/-^ flies. (A) Individual alpha/beta (left), alpha’/beta’ (middle), and gamma Kenyon cells (right) were photo-labeled in *Orco*^+/+^ (top) and *Orco*^-/-^ (bottom) flies, and the post-synaptic terminals formed by these neurons in the mushroom body calyx — called ‘claws’ — were imaged. Scale bar is 15 μm. (B-D) The total number of dendritic branches (B) and claws (C) were counted, and the total branch length formed by a neuron was measured (D) in *Orco*^+/+^ (green) and *Orco*^-/-^ flies (blue); the long black bars represent the median whereas the short black bars represent the 25th and 75th percentile of the data; the asterisk indicates values that were statistically different (*p* < 0.05). The statistical significance, or ‘*p*-value’, was measured using the Mann-Whitney U test.

Altogether, these results suggest that both projection neurons and Kenyon cells formed in *Orco*^-/-^ flies show no obvious morphological defects.

### Mushroom body input connections in *Orco*^-/-^ flies are biased and random

If sensory activity affects the way projection neurons connect to Kenyon cells, we would expect these connections to be qualitatively and quantitatively different in *Orco*^+/+^ and *Orco*^-/-^ flies. To compare global and more subtle connectivity patterns between genotypes, we used a neuronal tracing technique we previously devised [9]. In short, individual Kenyon cells were photo-labeled, and their input projection neurons were identified using dye electroporation. With this technique the inputs of hundreds of Kenyon cells can be identified and reported in a connectivity matrix. Statistical analyses of the resulting matrix can be used to reveal patterns of connectivity, including randomness and biases. We generated two connectivity matrices using *Orco*^+/+^ and *Orco*^-/-^ flies — henceforth referred to as the ‘*Orco*^+/+^ matrix’ and the ‘*Orco*^-/-^ matrix’ — by mapping the inputs of 200 Kenyon cells in each genotype; each matrix contains 745 and 734 connections, respectively (Figure 5A).

**Figure 5.**
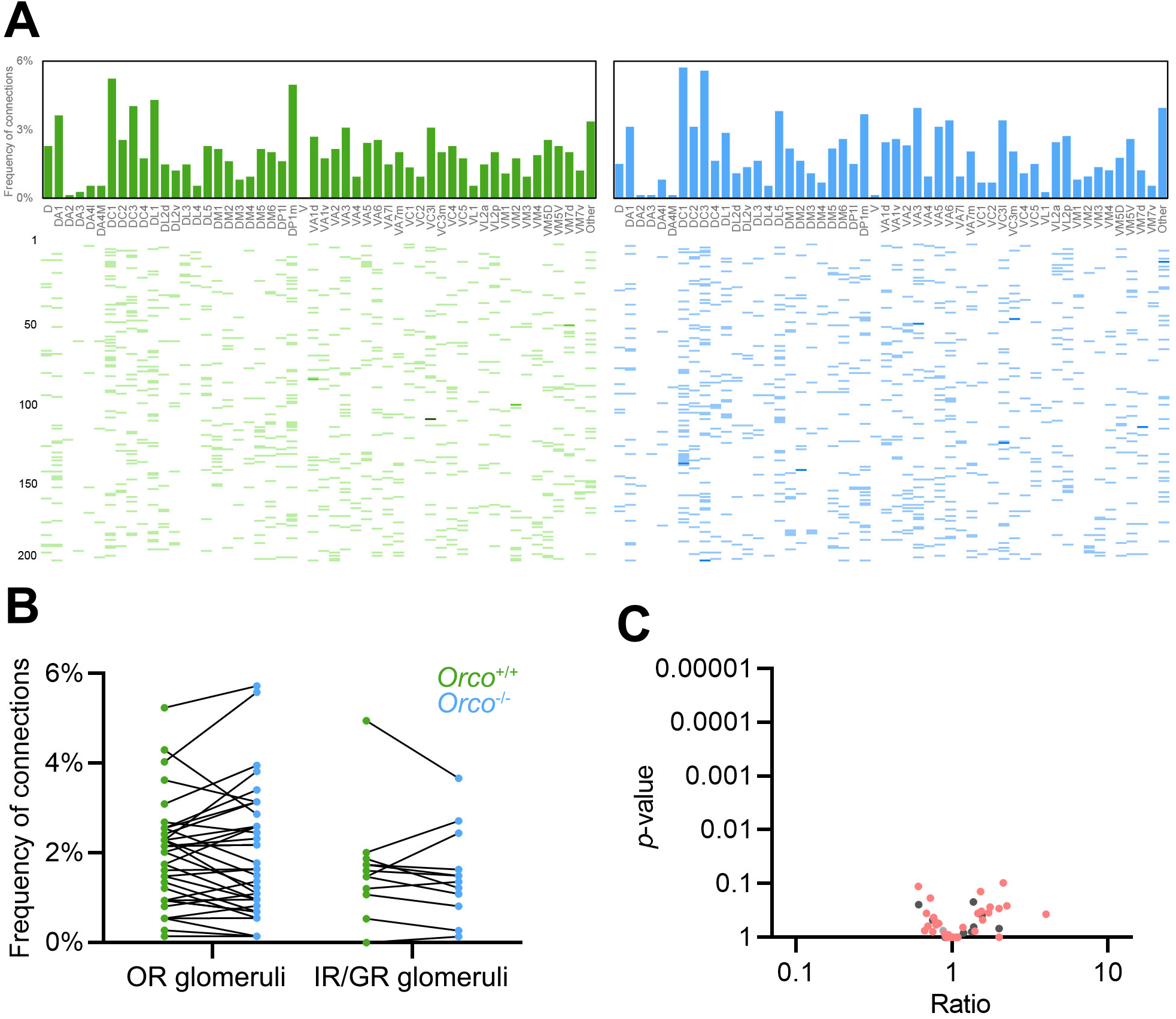
Connection frequencies are similar in *Orco*^+/+^ and *Orco*^-/-^ flies. (A) A total of 745 and 734 connections between projection neurons and Kenyon cells were mapped; all connections are reported in two connectivity matrices (*Orco*^+/+^: left panel and green; *Orco*^-/-^: right panel and blue). In each matrix, a row corresponds to a Kenyon cell (200 Kenyon cells per matrix) and each column corresponds to one of the 51 glomeruli; each colored bar indicates the input connections of a given Kenyon cell, and the intensity of the color indicates the number of connections found between a particular Kenyon cell and a given type of projection neuron (light: one connection; medium: two connections; dark: three connections). The bar graphs above the matrices represent the frequencies at which a particular type of projection neuron connects to Kenyon cells. (B) The frequencies at which different types of projection neuron connect to Kenyon cells in both data sets is shown (green: *Orco*^+/+^; blue: *Orco*^-/-^). Projection neurons are identified based on the glomeruli they innervate: ‘OR glomeruli’ refers to the projection neurons innervating glomeruli that receive input from OR-expressing neurons; ‘IR/GR glomeruli’ refers to projection neurons innervating glomeruli that receive input from IR/GR-expressing neurons. (B) The frequencies of connections measured for a given type of projection neuron in both genotypes are link with a black line. (C) The *p*-value measured for each glomerulus was plotted against the ratio of frequencies (ratio = frequency of connections in *Orco*^+/+^ / frequency of connections in *Orco*^-/-^) measured for each glomerulus (pink: projection neuron(s) receiving input from an OR glomerulus, dark grey: projection neurons receiving input from an IR or GR glomerulus; light grey: unknown). The statistical significance, or ‘*p*-value’, measured for each glomerulus was measured using the Fischer’s exact test; to control for false positives, *p*-values were adjusted with a false discovery rate using a Benjamini-Hochberg procedure. A ratio of 1 indicates that there is no shift in frequencies between the *Orco*^+/+^ and *Orco*^-/-^ flies whereas a ratio smaller than 1 indicates that a given type of projection neuron connects more frequently in *Orco*^-/-^ and a ratio greater than 1 indicates that a given type of projection neuron connects more frequently in *Orco*^+/+^.

We used different statistical analyses to compare these matrices. As a first step, we measured the frequencies at which projection neurons connect to Kenyon cells (Figure 5B,C and Figure S3). Lack of sensory activity could affect the frequencies at which projection neurons connect to Kenyon cells in at least two different ways. First, it is conceivable that the number of connections formed by projection neurons receiving input from ORs-expressing neurons would be higher in *Orco*^+/+^ flies, where they receive functional input, than in *Orco*^-/-^ flies, where their input neurons are silent. Such differences would be especially noticeable for projection neurons that connect to Kenyon cells at high frequencies in *Orco*^+/+^ flies, such as the DA1 projection neurons. However, we could not detect such differences: all projection neurons that receive input from ORs-expressing neurons — including the DA1 projection neurons — connect at similar frequencies in both genotypes (DA1 connectivity rates: *Orco*^+/+^: 3.62%, *Orco*^-/-^: 3.13%, *p*-value: 0.65; Figure 5B, C and Figure S3). Second, it is possible that the number of connections formed by projection neurons receiving input from IR/GRs-expressing neurons would be higher in *Orco*^-/-^ flies, where they are the only neurons that receive functional input, than in *Orco*^+/+^ flies. This would support the idea that projection neurons compete when connecting with Kenyon cells and that projection neurons that receive active input are advantaged. Such differences would be noticeable for projection neurons that connect to Kenyon cells at low frequencies in *Orco*^+/+^ flies, such as the VL1 projection neurons, as well as for projection neurons that connect at high frequencies, such as the DP1m neuron. However, we could not detect such differences: all projection neurons that receive input from IR/GR-expressing neurons — including the VL1 and DP1m projection neurons — connect at similar frequencies in both genotypes (VL1 connectivity rates: *Orco*^+/+^: 0.54%, *Orco*^-/-^: 0.27%, *p*-value: 0.68; DP1m: *Orco*^+/+^: 4.97%, *Orco*^-/-^: 3.68%, *p*-value: 0.22; Figure 5B, C and Figure S3). We could not detect shifts in connectivity frequencies that were significant; the most significant shift detected was for the VC4 projection neurons (VC4 connectivity rates: *Orco*^+/+^: 2.28%, *Orco*^-/-^: 1.09%, *p*-value: 0.097; Figure 5B, C and Figure S3). The frequencies measured for each projection neuron are largely similar across Kenyon cell types in both genotypes with the exception of the VL2p projection neurons that connect more frequently to alpha’/beta’ Kenyon cells in *Orco*^-/-^ flies than in *Orco*^+/+^ flies (VL2p connectivity rates: *Orco*^+/+^: 1.70%, *Orco*^-/-^: 11.76%, *p*-value: 0.0091; Figure S3). This difference is however most likely due to the small sample size: there are only 34 and 16 alpha’/beta’ Kenyon cells in the *Orco*^+/+^ and *Orco*^-/-^ matrices, respectively.

As a second step, we used the Jensen-Shannon distance — a statistical method that measures the likeness of two probability distributions — as a global readout of similarity in the observed connectivity biases. A distance of 0 would indicate that the two probability distributions are identical, whereas larger distances would indicate that the two probability distributions are different. To gauge the extent to which the Jensen-Shannon distance indicates likeness in our data sets, we generated two different shuffled versions of the connectivity matrices. In one version, called ‘shuffle’, the connections between projection neurons and Kenyon cells were randomly shuffled by choosing input projection neurons without replacement while keeping the number of Kenyon cells and number of projection neuron inputs to each Kenyon cell consistent with the experimental matrices. In the other version, called ‘fixed-shuffle’, the connections were randomly shuffled but the frequencies at which projection neurons connect to Kenyon cells were fixed to reflect the frequencies measured experimentally. When we compared each of the experimental matrices — the *Orco*^+/+^ and *Orco*^-/-^ matrices — to their shuffled versions, we obtained distances ranging from 0.224 to 0.266; when we compared the experimental matrices to their fixed-shuffle versions, we obtained distances ranging from 0.087 to 0.149 (Figure 6A). The Jensen-Shannon distance measured when comparing the *Orco*^+/+^ and *Orco*^-/-^ matrices is 0.115 — a value in the range of the values obtained with the fixed-shuffle matrices but outside the range of the values obtained with the shuffle matrices — suggesting that the distribution of connectivity frequencies is largely similar in both genotypes (Figure 6A).

**Figure 6.**
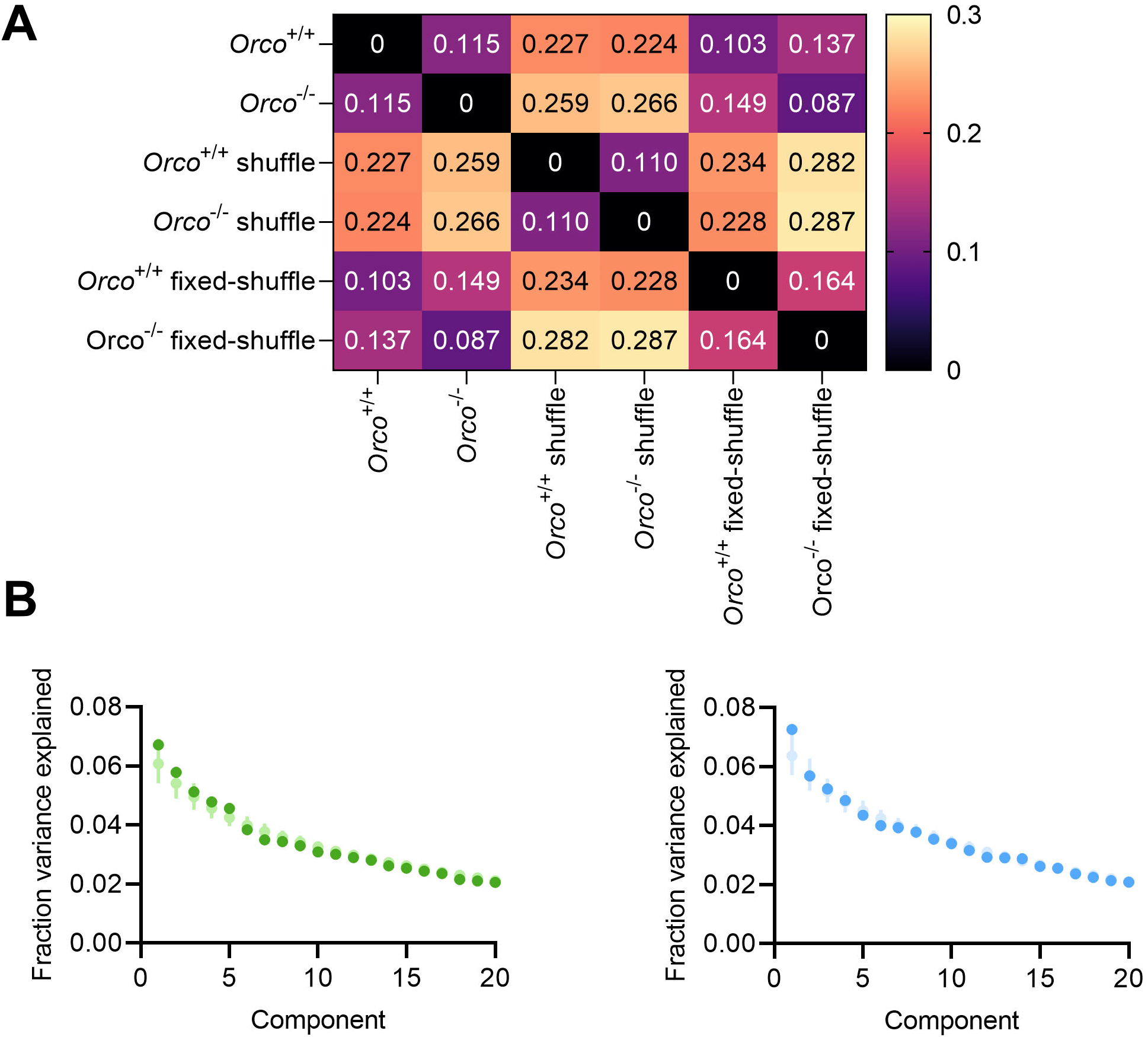
Mushroom body input connectivity is globally similar in *Orco*^+/+^ and *Orco*^-/-^ flies. (A) The Jensen-Shannon distances were measured between the experimental matrices, their shuffle version as well as their fixed-shuffle version. The color bar denotes the range in the distances measured. (B-C) Principal components were extracted using either the *Orco*^+/+^ (left panel, green) or the *Orco*^-/-^ (right panel, blue) connectivity matrices — using the experimental and fixed-shuffle versions — and the fraction of the variance explained by each component was measured (dark circles: experimental matrices, light circles: fixed-shuffle versions); error bars represent 95% confidence interval.

As a final step, we used an unbiased search for structural patterns that might exist in the connectivity matrices and that are not detectable by simply examining connectivity frequencies. To this end, we extracted correlations within each connectivity matrix — experimental and fixed-shuffle matrices — using principal component analysis (Figure 6B). The percent variance associated with the different principal component projections provides a sensitive measure of structure within each matrix [9]. For all components, the percent variance measured for the experimental matrix falls within the range measured for the fixed-shuffle matrices, suggesting that there are no structural patterns in the *Orco*^+/+^ and *Orco*^-/-^ matrices other than the biases in connectivity frequencies. It is worth noting that a recent study identified in the *Drosophila* hemibrain connectome a group of projection neurons that appear to preferentially connect to the same Kenyon cells but we could not find evidence for such group structure in our data sets (Figure S4) [27].

Altogether, these results show that the mechanisms underlying the biased and random connectivity pattern observed between projection neurons and Kenyon cells do not depend on sensory activity and are therefore most likely hardwired.

## DISCUSSION

In this study, we investigated whether the biased and random connectivity pattern observed between projection neurons and Kenyon cells forms depending on available sensory information. We first showed that in *Orco*^-/-^ flies, Kenyon cells fail to respond to odors detected by ORs but respond normally to odors detected by IRs. Second, we showed that — despite being partially anosmic — *Orco*^-/-^ flies develop a largely normal olfactory circuit: all glomeruli forming the antennal lobe can be identified in *Orco*^-/-^ flies, and the projection neurons and Kenyon cells found in *Orco*^-/-^ flies are morphologically similar to those found in *Orco*^+/+^ flies. Third, we mapped a large number of connections between projection neurons and Kenyon cells in *Orco*^+/+^ and *Orco*^-/-^ flies and compared the observed connectivity patterns using various statistical analyses. We could not detect any significant differences: projection neurons connect with Kenyon cells at similar frequencies in both data sets. Altogether these results suggest that the biased and random connectivity pattern observed between projection neurons and Kenyon cells forms independently of sensory activity.

It is possible that there are subtle connectivity patterns established by sensory activity that have eluded our analyses. For instance, a previous study identified a small number of alpha/beta and alpha‘-beta’ Kenyon cells that receive input more frequently from a group of ten glomeruli tuned to different food odors [27]. However, we failed to detect similar group structure in both the *Orco*^+/+^ and *Orco*^-/-^ connectivity matrices, suggesting that our mapping strategy cannot be used to reveal such subtle connectivity patterns. More exhaustive mapping strategies — such as the derivation of an *Orco*^-/-^ connectome — might be necessary to determine whether this possible group structure might result from sensory activity. However, our technique is clearly able to detect global, population-level connectivity patterns, and our results show that these patterns form independently of sensory activity.

This independence of sensory activity came as a surprise in light of the evidence suggesting that the synapse between projection neurons and Kenyon cells is plastic in *Drosophila melanogaster*. Previous study noticed that, when a few projection neurons are silenced or chronically activated, the volumes of their presynaptic terminals change in the mushroom body [28,29]. Another study showed that appetitive conditioning leads to an increase in the number of synapses formed between the projection neurons activated by the conditioned odor and Kenyon cells [30]; similar observations have been made in honeybees (review in [31]). However, none of these studies could determine whether these plastic changes lead to lasting changes in connectivity pattern. Our results partially support these findings: we found that lack of sensory activity leads to a small but significant increase in the size of the presynaptic terminals formed by projection neurons in the mushroom body. However, our results also demonstrate that these sensory-based changes have little to no effect on the global connectivity pattern between projection neurons and Kenyon cells.

Our results support the idea that the biased and random connectivity pattern observed between projection neurons and Kenyon cells results from hardwired mechanisms. It is possible that such mechanisms regulate spontaneous activity either at the level of olfactory sensory neurons or at the level of projection neurons. This possibility appears, however, unlikely considering that olfactory sensory neurons in *Orco*^-/-^ flies show drastically reduced levels of spontaneous activity [17]. Likewise, a previous study showed the DL1 and VM2 projection neurons in flies lacking their cognate receptor genes — *OR10a*^-/-^ and *OR43b*^-/-^ flies, respectively — are virtually silent and display low to no spontaneous activity [22]. Thus, the hardwired mechanisms leading to biases in connectivity most likely involve synapse promoting factors which may be differentially expressed in overrepresented *versus* underrepresented neurons. A recent study found that the number of Kenyon cells affects the number of presynaptic sites formed by projection neurons: the more Kenyon cells there are, the fewer presynaptic sites [32]. This result suggests that Kenyon cells might release a synapse-promoting signal that is differentially detected by projection neurons leading to the observed connectivity biases.

In theory, biased and random input connections are connectivity patterns that antagonize each other: the lack of structure afforded by randomization of inputs enables the mushroom body to represent olfactory information with as many unique activity patterns as possible, whereas the structure imposed by biases skews these representations to prioritize a few ethologically meaningful odors. Our finding that biases do not simply reflect the concrete chemosensory ecology of a fly but are hardwired suggests that this connectivity pattern has been shaped on a long-term evolutionary timescale. It is tempting to speculate that biases might prepare the mushroom body to learn more efficiently from the chemosensory environment present in the particular ecological niche of a species. This finding has important ramifications for our understanding of how such fairly subtle, yet significant connectivity patterns develop and evolve as well as our understanding of how biases in connectivity might be evolutionarily adaptive.

## AKNOWLEDGMENTS

We thank members of the Caron laboratory for comments on the manuscript; Anita Devineni for sharing the code used to analyze imaging data; Cody Orton for the initial characterization of Kenyon cells; Adam Lin for preparation of the standard cornmeal agar medium; Ashley Platt for assistance with general laboratory concerns. This work has been funded by grants from the National Institute for Neurological Disorders and Stroke (R01 NS 106018, R01 NS 1079790 and R01 EB 029858), the National Science Foundation (IOS 2042397 and DBI 1707398) and the Gatsby Charitable Foundation. Further financial support was provided by the DOE CSGF (DE-SC0022158) (I.G.), the Research Scholar Award (M.S.J.), the University Research Opportunities Program (M.S.J.), the Burroughs Wellcome Foundation (A.L.K.), the McKnight Endowment Fund (A.L.K.), the Simons Collaboration on the Global Brain (A.L.K.) and the Georges S. and Dolores Eccles Foundation (S.J.C.C.).

## AUTHORS CONTRIBUTIONS

T.H. and S.J.C.C. conceived the project. A.J.M. collected and analyzed the calcium imaging data with the help of T.H.; H.M.S. generated and analyzed the antennal lobe reconstructions; T.H. photo-labeled projection neurons and Kenyon cells and analyzed their morphology with the help of M.S.J.; T.H. generated the connectivity matrices; I.G. and A.L.K. performed the statistical analyses of the connectivity matrices. T.H. and S.J.C.C. wrote the manuscript with input from all other authors.

---

**Table S1.** Volumes of the glomeruli reconstructed in *Orco*^+/+^ and *Orco*^-/-^ flies. Individual glomeruli were reconstructed and their volumes was measured in five different samples for each genotype. The statistical significance, or ‘*p*-value’, was measured using the Mann-Whitney U test.

| Glomerulus | Type of receptor in sensory neurons | Adjusted *p*-value | Mean ± s.d. volume in *Orco*<sup>+/+</sup> flies ($\mu\text{m}^3$) | Mean ± s.d. volume in *Orco*<sup>-/-</sup> flies ($\mu\text{m}^3$) |
| --- | --- | --- | --- | --- |
| D | OR | 0.101190 | 1578 ± 226 | 997 ± 270 |
| DA1 | OR | 0.101190 | 3994 ± 370 | 3377 ± 252 |
| DA2 | OR | 0.067460 | 1040 ± 96 | 1462 ± 189 |
| DA3 | OR | 0.765528 | 374 ± 49 | 352 ± 86 |
| DA4l | OR | 0.765528 | 768 ± 60 | 747 ± 71 |
| DA4m | OR | 0.858095 | 707 ± 88 | 693 ± 167 |
| DC1 | OR | 0.765528 | 1628 ± 98 | 1596 ± 246 |
| DC2 | OR | 0.089947 | 1223 ± 74 | 1011 ± 87 |
| DC3 | OR | 0.464286 | 1527 ± 162 | 1453 ± 58 |
| DL1 | OR | 0.089947 | 1522 ± 139 | 1269 ± 75 |
| DL3 | OR | 0.141667 | 1193 ± 110 | 909 ± 201 |
| DL4 | OR | 0.067460 | 419 ± 25 | 279 ± 28 |
| DL5 | OR | 0.607143 | 1452 ± 166 | 1616 ± 254 |
| DM1 | OR | 0.231293 | 3495 ± 783 | 2617 ± 288 |
| DM2 | OR | 0.377778 | 1763 ± 206 | 1928 ± 139 |
| DM3 | OR | 0.101190 | 944 ± 37 | 830 ± 55 |
| DM4 | OR | 0.101190 | 1772 ± 351 | 1328 ± 163 |
| DM5 | OR | 0.141667 | 1229 ± 147 | 960 ± 134 |
| DM6 | OR | 0.307619 | 1206 ± 66 | 1368 ± 157 |
| VA1d | OR | 0.464286 | 2818 ± 46 | 2656 ± 355 |
| VA1v | OR | 0.579794 | 3984 ± 238 | 3796 ± 451 |
| VA2 | OR | 0.101190 | 3363 ± 265 | 2805 ± 221 |
| VA3 | OR | 0.101190 | 1796 ± 122 | 1453 ± 144 |
| VA4 | OR | 0.765528 | 957 ± 41 | 985 ± 175 |
| VA5 | OR | 0.765528 | 1033 ± 137 | 1130 ± 174 |
| VA6 | OR | >0.999999 | 2237 ± 162 | 2188 ± 372 |
| VA7l | OR | 0.067460 | 911 ± 65 | 723 ± 51 |
| VA7m | OR | 0.307619 | 1092 ± 33 | 990 ± 110 |
| VC1 | OR | 0.067460 | 1210 ± 124 | 941 ± 71 |
| VC2 | OR | 0.141667 | 982 ± 91 | 835 ± 54 |
| VC4 | OR | 0.101190 | 1061 ± 208 | 808 ± 82 |
| VM2 | OR | 0.765528 | 1066 ± 140 | 1046 ± 42 |
| VM3 | OR | 0.307619 | 1302 ± 38 | 1235 ± 230 |
| VM5d | OR | 0.377778 | 1159 ± 210 | 973 ± 206 |
| VM5v | OR | 0.141667 | 1429 ± 268 | 981 ± 228 |
| VM7d | OR | 0.307619 | 1458 ± 306 | 1064 ± 290 |
| VM7v | OR | 0.464286 | 1674 ± 296 | 1432 ± 182 |
| DC4 | IR | 0.464286 | 1290 ± 223 | 1081 ± 258 |
| DL2d | IR | 0.089947 | 1626 ± 265 | 2230 ± 214 |
| DL2v | IR | 0.377778 | 1591 ± 176 | 1747 ± 130 |
| DP1l | IR | 0.377778 | 2594 ± 241 | 2736 ± 240 |
| DP1m | IR | 0.067460 | 3969 ± 404 | 5437 ± 247 |
| VC5 | IR | 0.858095 | 1316 ± 127 | 1311 ± 204 |
| VL1 | IR | 0.377778 | 3171 ± 229 | 3487 ± 334 |
| VL2a | IR | 0.579794 | 2351 ± 187 | 2506 ± 245 |
| VL2p | IR | 0.765528 | 2727 ± 196 | 2747 ± 270 |
| VM1 | IR | 0.067460 | 905 ± 91 | 1105 ± 97 |
| VM4 | IR | 0.858095 | 1126 ± 134 | 1077 ± 188 |
| V | GR | 0.858095 | 2461 ± 333 | 2525 ± 258 |
| VC3l | Unknown/mixed | 0.579794 | 1143 ± 289 | 980 ± 234 |
| VC3m | Unknown/mixed | 0.765528 | 1042 ± 109 | 1009 ± 123 |

**Figure S1.**
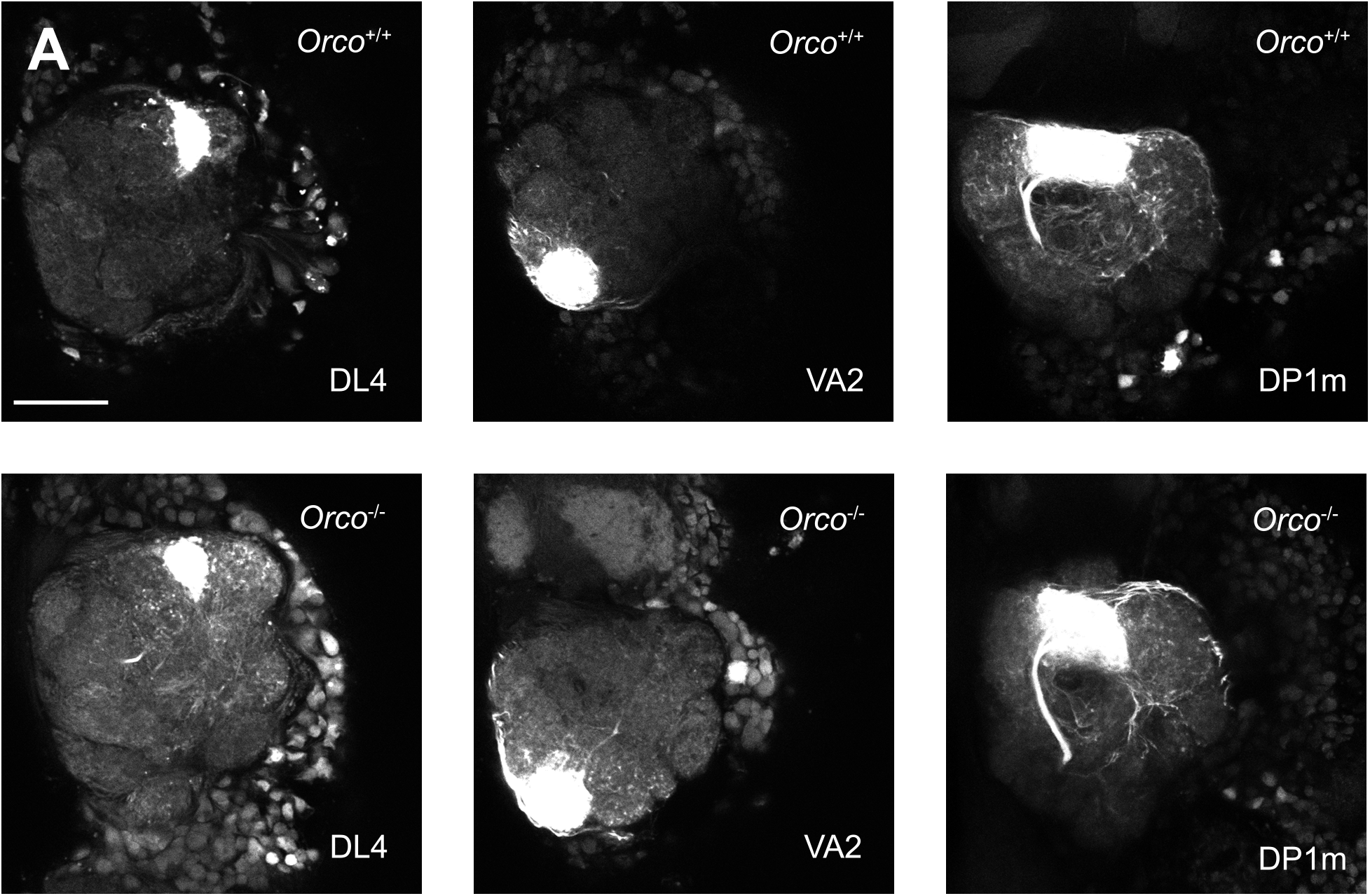
Photo-labeled glomeruli. The DL4 (left), VA2 (center), DP1m (right) glomeruli were photo-labeled in *Orco*^+/+^ (top) and *Orco*^-/-^ (bottom) flies and imaged. Scale bar is 15 μm.

**Figure S2.**
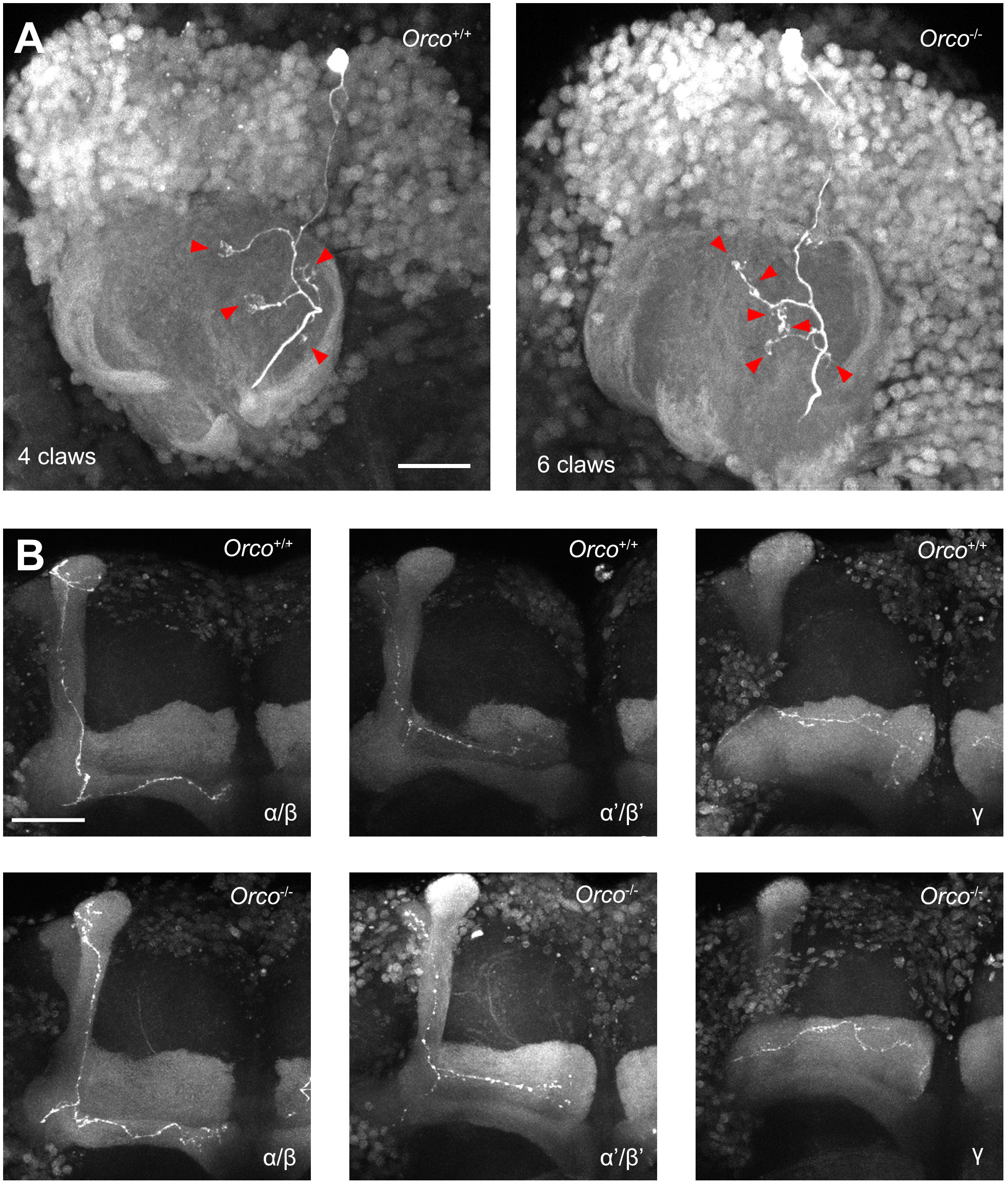
Photo-labeled Kenyon cells. (A-B) Individual Kenyon cells were photo-labeled in *Orco*^+/+^ and *Orco*^-/-^ flies. (A) The number of claws formed by a Kenyon cell was counted; each claw is demarcated by a red arrow. Here is an example of an alpha/beta Kenyon cell found in an *Orco*^+/+^ fly that has four claws (left) and an alpha’/beta’ Kenyon cell found in an *Orco*^-/-^ fly that has six claws (right). Scale bar is 15 μm. (B) The type of Kenyon cell that was photo-labeled was identified based on its axonal innervation pattern in the mushroom body lobe. Here are examples of an alpha/beta (left panels), alpha’/beta’ (middle panels) and gamma (right panels) Kenyon cells found in *Orco*^+/+^ (top) and *Orco*^-/-^ flies (bottom). Scale bar is 30 μm.

**Figure S3.**
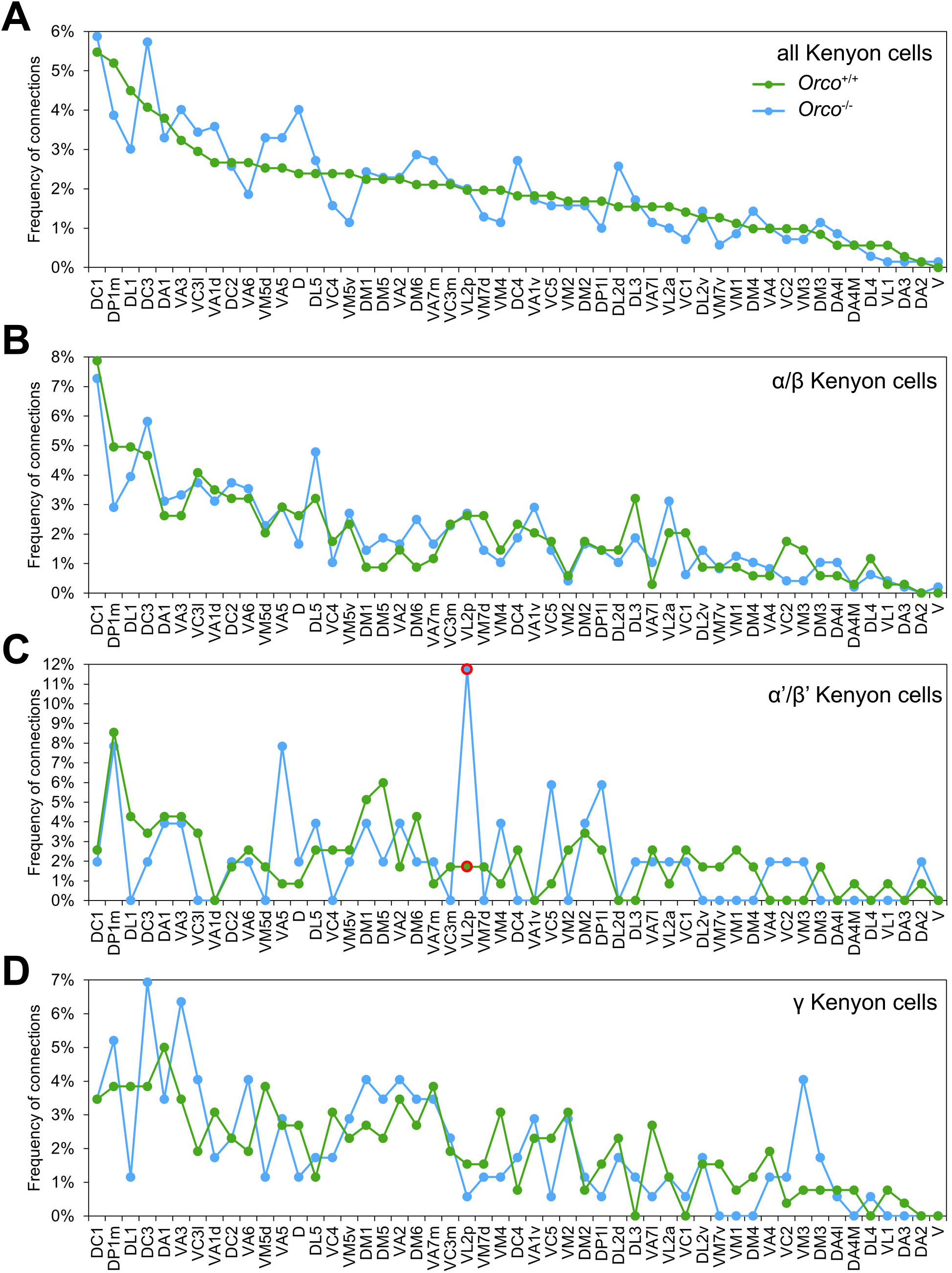
Distributions of connectivity frequencies. The distributions of connectivity frequencies obtained in the experimental datasets — *Orco*^+/+^ (green) and *Orco*^-/-^ (blue) — were plotted and compared across all Kenyon cells (top), alpha/beta Kenyon cells (middle top), alpha’/beta’ Kenyon cells (middle bottom) and gamma Kenyon cells (bottom). The statistical significance, or ‘*p*-value’ measured for each glomerulus was measured using the Fischer’s exact test; red outlines indicate values that were statistically different (*p* < 0.01).

**Figure S4.**
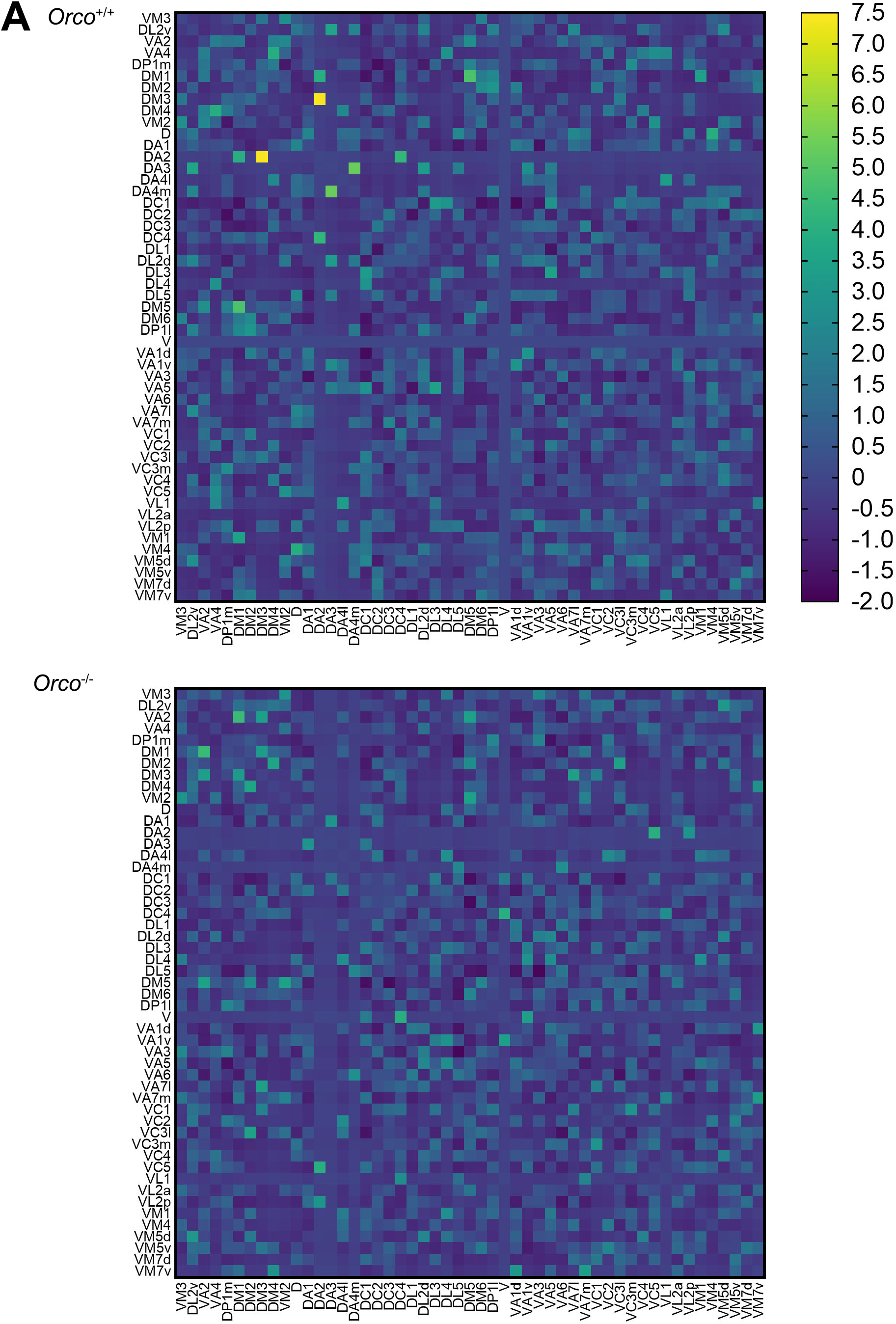
Conditional input matrices. In each conditional input matrix, each cell indicates whether a Kenyon cell is more, equally or less likely than chance to receive an input from a type of projection neuron (column) given that this Kenyon cell receives an input from another type of projection neuron (row). The color bar denotes the frequency measured for each pair. The projection neurons found to connect preferentially to the same Kenyon cells in Zheng et al. 2020 are listed first in each row and column; the remaining projection neurons are listed in alphabetical order.

## MATERIALS AND METHODS

### Fly stocks

Flies (*Drosophila melanogaster*) were fed on standard cornmeal agar medium and raised in a controlled environmental chamber (Percival Scientific, Inc.) that maintains a temperature of 25°C and 60% humidity under a 12 hours light-12 hours dark cycle. Crosses were set up and reared under the same conditions, but the standard cornmeal agar medium was supplemented with dry yeast. The strains used in this study are described in the table below.

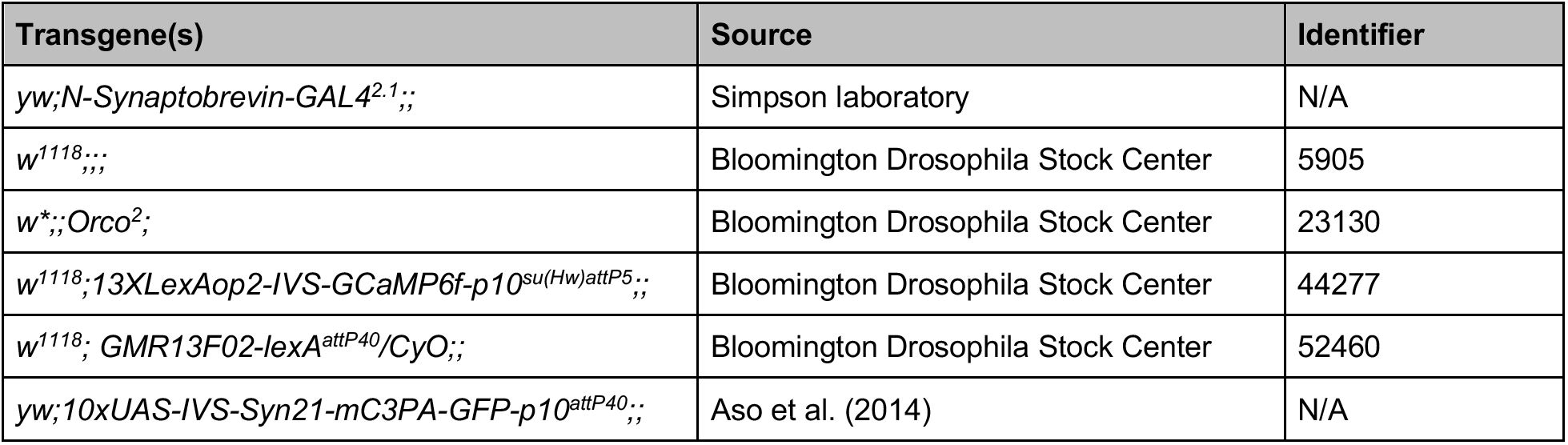

In all experiments, three-day old females were used. For the functional imaging experiments, we used the following transgenic lines: *w; [R13F02-LexA]^attP40^,[LexAop-GCaMP6f]^su(Hw)attP5^*/+;; (referred to in the text as ‘*Orco*^+/+^’) and *w; [R13F02-LexA]^attP40^, [LexAop-GCaMP6f]^su(Hw)attP5^/+;Orco^2^*; (referred to in the text as *Orco*^-/-^’). For the antennal lobe reconstructions, we used the following transgenic lines: *w*;+;+;+, (referred to in the text as ‘*Orco*^+/+^’) *w;+;Orco^2^*;+ (referred to in the text as *Orco*^-/-^’). For the photolabeling and connectivity mapping experiments, we used the following transgenic lines: *w;[N-Synaptobrevin-GAL4-GAL4]^2.1^,[10xUAS-IVS-Syn21-mC3PA-GFP-p10]^attP40^*;; (referred to in the text as ‘*Orco*^+/+^’) and *w;[N-Synaptobrevin-GAL4-GAL4]^2.1^,[10xUAS-IVS-Syn21-mC3PA-GFP-p10]^attP40^;Orco^2^*; (referred to in the text as ‘*Orco*^-/-^’). All *Orco*^-/-^ stocks were genotyped every other week by performing PCR using a previously established protocol [17].

### Functional imaging

Calcium imaging experiments were performed on immobilized flies. Flies were immobilized underneath an imaging chamber — a platform with an opening attached to a reservoir of saline — using a combination of clear tape (Shurtape Technologies, DUC280068) and UV glue (Bondic, SK8024). A hole was cut in the cuticle of the fly, above the mushroom body, and the exposed brain was submerged in saline (108 mM NaCl, 5 mM KCl, 5 mM HEPES, 5 mM Trehalose, 10 mM Sucrose, 1 mM NaH_2_PO_4_, 4 mM NaHCO_3_, 2 mM CaCl_2_, 4 mM MgCl_2_, pH≈7.3). Immobilized flies were exposed to two odors — isopentyl acetate (Sigma-Aldrich, 112674) diluted 5% volume to volume in paraffin oil (Fluka Analytical, 76235) and acetic acid (Fisher Scientific, A385) diluted in 5% volume to volume in water — using a stimulus controller (Ockenfels Syntech, CS 55). For imaging calcium transients, we used a two-photon laser scanning microscope (Bruker, Ultima Investigator) equipped with an ultrafast Chameleon Ti:Saphirre laser (Coherent) modulated by Pockels Cells (Conotopics). The laser power used for each experiment varied from 14 to 24 mW. Calcium transients were recorded in the calyx of the mushroom body using the following sequence: 5 seconds odor ‘off’, 2 seconds odor ‘on’ and 8 seconds odor ‘off’. This sequence was repeated five times. Image sequences were collected with either a galvo and 512 by 512 pixels resolution with 0.8μs dwell time and 1.64 fps (for generating the heatmaps shown in Figure 1A) or a resonant galvo and 512 by 512 resolution with 15.081 fps (for generating the traces shown in Figure 1B,C). Calcium transients were analyzed using a custom MATLAB code based on previously published codes [33–35]. To correct for movement, images were registered within and between trials using a sub-pixel registration algorithm [36]. Heatmaps were generated by averaging the intensity of individual pixels (F_0_: The entire five seconds of the first off-period combined with the last two seconds of the second off-period; F: The entire two seconds of the on-period). Traces were generated by averaging the calcium transients detected in the main calyx (F_0_: The entire five seconds of the first off-period combined with the last two seconds of the second off-period; F: The entire two seconds of the on-period).

### Reconstructing antennal lobes

Antennal lobes were reconstructed from confocal images of immuno-stained brains. The brains of flies were dissected at room temperature in a phosphate buffered saline solution or PBS (Sigma-Aldrich, P5493), fixed in 2% paraformaldehyde (Electron Microscopy Sciences, 15710) for 45 minutes at room temperature, washed five times in PBST (PBS with 1x Triton, Sigma-Aldrich, T8787) at room temperature, blocked with 5% goat Serum (Jackson ImmunoResearch Laboratories) in PBST for 30 minutes at room temperature, and incubated in a solution that contained the primary antibody (1:20 in 5% Goat Serum/PBST, Developmental Studies Hybridoma Bank, nc82, AB_2314866) at 4°C overnight. On the following day, brains were washed four times in PBST and incubated in a solution that contained the secondary antibody (1:500 in 5% Goat Serum/PBST, Thermal Fisher, goat anti-mouse Alexa Fluor 488, AB_2576217) at 4°C overnight. On the following day, brains were washed four times in PBST and mounted on a slide (Fisher Scientific, 12-550-143) using the mounting media VECTASHIELD (Vector Laboratories Inc., H-1000). Immuno-stained brains were imaged using an LSM 880 confocal microscope (Zeiss). Each antennal lobe was reconstructed from a confocal image using the segmentation software Amira (FEI Visualization Sciences Group, version 2020.3.1). Individual glomeruli were reconstructed via manual segmentation: boundaries were demarcated by hand and interpolated. Glomeruli were assigned identities according to their position based on the available anatomical maps and the *Drosophila melanogaster* hemibrain connectome v1.2.1 [4,5,26,37]. Glomerular volumes were calculated from the reconstructed voxel size, and the sum of those volumes were used to calculate whole antennal lobe volumes.

### Photo-labeling projection neurons and Kenyon cells

Neurons were photo-labeled based on a previously published protocol [38]. In short, brains were dissected in saline (108 mM NaCl, 5 mM KCl, 5 mM HEPES, 5 mM Trehalose, 10 mM Sucrose, 1 mM NaH_2_PO_4_, 4 mM NaHCO_3_, 2 mM CaCl_2_, 4 mM MgCl_2_, pH≈7.3), treated for 1 minute with 2 mg/ml collagenase (Sigma-Aldrich) and mounted on a piece of Sylgard placed at the bottom of a Petri dish. Each brain was either mounted with its anterior side facing upward (for photo-labeling projection neurons) or with its posterior side facing upward (for photo-labeling Kenyon cells). The photo-labeling and image acquisition steps were performed using a two-photon laser scanning microscope (Bruker, Ultima) with an ultrafast Chameleon Ti:Sapphire laser (Coherent) modulated by Pockels Cells (Conotopics). During the photo-labeling step, the laser was tuned to 710 nm and about 5 to 30 mW of laser power was used; during the image acquisition step, the laser was tuned to 925 nm and about 1 to 14 mW of laser power was used. Both power values were measured behind the objective lens. A 60X water-immersion objective lens (Olympus) was used for both photo-labeling and image acquisition. A GaAsP detector (Hamamatsu Photonics) and PMT detector (Bruker) were used for measuring green and red fluorescence, respectively. Photolabeling was performed by drawing a region of interest — on average 1.0 × 1.0 μm — either in the center of the targeted glomerulus (for labeling projection neurons) or in the center of the soma (for labeling Kenyon cells); each pixel was scanned 8 times. Image acquisition was performed at a resolution of 512 by 512 pixels with a pixel size of 0.39 μm and a pixel dwell time of 4 μs; each pixel was scanned 2 times.

### Mapping Kenyon cell input connections using dye electroporation

The projection neurons connecting to a photo-labeled Kenyon cell were identified as described before [9] with some modification. In short, electrodes were made by pulling borosilicate glass pipette with filament (Sutter Instruments, BF100-50-10) to a resistance of 9-11 MΩ, fire-polished using a micro-forge (Narishige) to narrow their opening, and backfilled with 100mg/ml 3000-Da Texas-dextran dye (Thermo-Fisher, D3328). Under the guidance of a two-photon microscope (Bruker, Ultima), an electrode was centered into the post-synaptic terminal — or ‘claw’ — of a photo-labeled Kenyon cell using a motorized micromanipulator (Sutter Instruments). Short current pulses (each 10-50 V in amplitude and 0.5 millisecond long) were applied until the projection neuron connecting to the targeted Kenyon cell claw was visible. Not all the projection neurons connecting to a given Kenyon cells were dye-filled but on average 4 ± 1 of the claws formed by a given Kenyon cell were dye-filled. An image of the antennal lobe was acquired at the end of the procedure. Dye-labeled glomeruli were identified based on their shape, position and the location of their soma as defined in the available anatomical maps and the *Drosophila melanogaster* hemibrain connectome v1.2.1 [4,5,26,37].

### Quantifying morphological features

All quantifications were done blindly without prior knowledge of the genotype of a sample. Representative images of antennal lobes, projection neurons, Kenyon cells were projected at maximal intensity using the ImageJ/Fiji software (National Institutes of Health [39]). The multipoint tool was used to count the number of projection neurons, bouton clusters and primary branches as well as the number of Kenyon cell claws. Projection neurons were counted based on the number of photo-labeled bodies observed in the anterior or lateral clusters of the antennal lobe. Bouton clusters were defined as aggregates decorating individual dendritic branches, and each bouton cluster is at least 1μm apart from another. Primary branches were defined as processes that emerge from the main axonal projection traversing the calyx of the mushroom body. Claws were defined as cup-like endings located at the end of a dendritic process formed by a Kenyon cell. The total bouton volume was measured using Fluorender (University of Utah Scientific Computing and Imaging Institute; version 2.26.2 [40,41]). Boutons were traced using the paint brush function. To efficiently distinguish boutons from the background, the ‘Edge Detect’ parameter was kept on and the ‘Edge STR’ was fixed at 0.505, while the selection threshold was adjusted to different values depending on signal intensity. We reported the ‘Physical Size’ value as total bouton volume. The length of the branches and claws formed by a Kenyon cell was quantified using the ‘Simple Neurite Tracer’ plugin and the ImageJ/Fiji software (National Institutes of Health [39,42]). Branches and claws were traced and the resulting trace was measured using the ‘Measure’ function. We reported the value measured using the ‘Cable Length’ function as the length.

### Statistical analyses

Statistical analyses for the data shown in Figure 1 to 4: *p*-values were computed using the Mann-Whitney U test. Statistical significance is indicated as *p* < 0.05 (*), *p* < 0.01 (**) and *p* < 0.001 (***). Statistical analyses for the data shown in Figure 5C: *p*-values were computed using the Fischer’s exact test. To control for false positives, *p*-values were adjusted with a false discovery rate of 10% using a Benjamini-Hochberg procedure. Statistical analyses for the data shown in Figure S4: The methods used to generate the conditional input matrices have been described in a previous study [27]. In short, each cell in the conditional input matrices indicates whether a Kenyon cell is more, equally or less likely than chance to receive an input from a type of projection neuron (column) given that this Kenyon cell receives an input from another type of projection neuron (row). Each observed projection neuron–Kenyon cell connection is treated as a single count. The observed number of counts for a given pair of neurons is compared to the distribution of counts generated using a null model. In the null model, 1,000 connectivity matrices were generated by randomly shuffling the connections recorded in the corresponding experimental matrix while keeping the number of input each Kenyon cell receives and the frequencies at which projection neurons connect to Kenyon cells constant; these matrices are referred in the main text as ‘fixed shuffle matrices’. For each pair of projection neurons, a z-score representing the number of standard deviations from the mean of the null distribution and the observed counts was computed. Unsupervised K-means clustering of the z-score matrix was used to group matrix entries. A K-means clustering (K=4) was applied to the projection neurons with similar z-scores.

## REFERENCES

1. Petrovic, M., and Schmucker, D. (2015). Axonal wiring in neural development: Target independent mechanisms help to establish precision and complexity. BioEssays 37, 996–1004.

2. Aso, Y., Hattori, D., Yu, Y., Johnston, R.M., Iyer, N.A., Ngo, T.-T.B., Dionne, H., Abbott, L.F., Axel, R., Tanimoto, H., et al. (2014). The neuronal architecture of the mushroom body provides a logic for associative learning. eLife 3, e04577.

3. Li, F., Lindsey, J.W., Marin, E.C., Otto, N., Dreher, M., Dempsey, G., Stark, I., Bates, A.S., Pleijzier, M.W., Schlegel, P., et al. (2020). The connectome of the adult Drosophila mushroom body provides insights into function. eLife 9, e62576.

4. Couto, A., Alenius, M., and Dickson, B.J. (2005). Molecular, Anatomical, and Functional Organization of the Drosophila Olfactory System. Curr. Biol. 15, 1535–1547.

5. Fishilevich, E., and Vosshall, L.B. (2005). Genetic and Functional Subdivision of the Drosophila Antennal Lobe. Curr. Biol. 15, 1548–1553.

6. Bates, A.S., Schlegel, P., Roberts, R.J.V., Drummond, N., Tamimi, I.F.M., Turnbull, R., Zhao, X., Marin, E.C., Popovici, P.D., Dhawan, S., et al. (2020). Complete Connectomic Reconstruction of Olfactory Projection Neurons in the Fly Brain. Curr. Biol. 30, 3183–3199.e6.

7. Schlegel, P., Bates, A.S., Stürner, T., Jagannathan, S.R., Drummond, N., Hsu, J., Capdevila, L.S., Javier, A., Marin, E.C., Barth-Maron, A., et al. (2021). Information flow, cell types and stereotypy in a full olfactory connectome. eLife 10, e66018.

8. Murthy, M., Fiete, I., and Laurent, G. (2008). Testing Odor Response Stereotypy in the Drosophila Mushroom Body. Neuron 59, 1009–1023.

9. Caron, S.J.C., Ruta, V., Abbott, L.F., and Axel, R. (2013). Random convergence of olfactory inputs in the Drosophila mushroom body. Nature 497, 113–7.

10. Gruntman, E., and Turner, G.C. (2013). Integration of the olfactory code across dendritic claws of single mushroom body neurons. Nat. Neurosci. 16, 1821–1829.

11. Litwin-Kumar, A., Harris, K.D., Axel, R., Sompolinsky, H., and Abbott, L.F. (2017). Optimal Degrees of Synaptic Connectivity. Neuron 93, 1153–1164.e7.

12. Zavitz, D., Amematsro, E.A., Borisyuk, A., and Caron, S.J.C. (2021). Connectivity patterns that shape olfactory representation in a mushroom body network model. bioRxiv, 2021.02.10.430647.

13. Kurtovic, A., Widmer, A., and Dickson, B.J. (2007). A single class of olfactory neurons mediates behavioural responses to a Drosophila sex pheromone. Nature 446, 542–546.

14. Silbering, A.F., Rytz, R., Grosjean, Y., Abuin, L., Ramdya, P., Jefferis, G.S.X.E., and Benton, R. (2011). Complementary Function and Integrated Wiring of the Evolutionarily Distinct Drosophila Olfactory Subsystems. J. Neurosci. 31, 13357–13375.

15. Ebrahim, S.A.M., Dweck, H.K.M., Stökl, J., Hofferberth, J.E., Trona, F., Weniger, K., Rybak, J., Seki, Y., Stensmyr, M.C., Sachse, S., et al. (2015). Drosophila Avoids Parasitoids by Sensing Their Semiochemicals via a Dedicated Olfactory Circuit. PLOS Biol. 13, e1002318.

16. Stensmyr, M.C., Dweck, H.K.M., Farhan, A., Ibba, I., Strutz, A., Mukunda, L., Linz, J., Grabe, V., Steck, K., Lavista-Llanos, S., et al. (2012). A Conserved Dedicated Olfactory Circuit for Detecting Harmful Microbes in Drosophila. Cell 151, 1345–1357.

17. Larsson, M.C., Domingos, A.I., Jones, W.D., Chiappe, M.E., Amrein, H., and Vosshall, L.B. (2004). Or83b Encodes a Broadly Expressed Odorant Receptor Essential for Drosophila Olfaction. Neuron 43, 703–714.

18. Benton, R., Sachse, S., Michnick, S.W., and Vosshall, L.B. (2006). Atypical Membrane Topology and Heteromeric Function of Drosophila Odorant Receptors In Vivo. PLoS Biol. 4, e20.

19. Jones, W.D., Cayirlioglu, P., Kadow, I.G., and Vosshall, L.B. (2007). Two chemosensory receptors together mediate carbon dioxide detection in Drosophila. Nature 445, 86–90.

20. Kvon, E.Z., Kazmar, T., Stampfel, G., Yáñez-Cuna, J.O., Pagani, M., Schernhuber, K., Dickson, B.J., and Stark, A. (2014). Genome-scale functional characterization of Drosophila developmental enhancers in vivo. Nature 512, 91–95.

21. Berdnik, D., Chihara, T., Couto, A., and Luo, L. (2006). Wiring Stability of the Adult Drosophila Olfactory Circuit after Lesion. J. Neurosci. 26, 3367–3376.

22. Olsen, S.R., Bhandawat, V., and Wilson, R.I. (2007). Excitatory Interactions between Olfactory Processing Channels in the Drosophila Antennal Lobe. Neuron 54, 89–103.

23. Chiang, A., Priya, R., Ramaswami, M., VijayRaghavan, K., and Rodrigues, V. (2009). Neuronal activity and Wnt signaling act through Gsk3-β to regulate axonal integrity in mature Drosophila olfactory sensory neurons. Development 136, 1273–1282.

24. Trible, W., Olivos-Cisneros, L., McKenzie, S.K., Saragosti, J., Chang, N.-C., Matthews, B.J., Oxley, P.R., and Kronauer, D.J.C. (2017). orco Mutagenesis Causes Loss of Antennal Lobe Glomeruli and Impaired Social Behavior in Ants. Cell 170, 727–735.e10.

25. Ryba, A.R., McKenzie, S.K., Olivos-Cisneros, L., Clowney, E.J., Pires, P.M., and Kronauer, D.J.C. (2020). Comparative Development of the Ant Chemosensory System. Curr. Biol. 30, 3223–3230.e4.

26. Scheffer, L.K., Xu, C.S., Januszewski, M., Lu, Z., Takemura, S., Hayworth, K.J., Huang, G.B., Shinomiya, K., Maitlin-Shepard, J., Berg, S., et al. (2020). A connectome and analysis of the adult Drosophila central brain. eLife 9, e57443.

27. Zheng, Z., Li, F., Fisher, C., Ali, I.J., Sharifi, N., Calle-Schuler, S., Hsu, J., Masoodpanah, N., Kmecova, L., Kazimiers, T., et al. (2020). Structured sampling of olfactory input by the fly mushroom body. bioRxiv, 2020.04.17.047167.

28. Kremer, M.C., Christiansen, F., Leiss, F., Paehler, M., Knapek, S., Andlauer, T.F.M., Förstner, F., Kloppenburg, P., Sigrist, S.J., and Tavosanis, G. (2010). Structural longterm changes at mushroom body input synapses. Curr. Biol. 20, 1938–44.

29. Doll, C.A., Vita, D.J., and Broadie, K. (2017). Fragile X Mental Retardation Protein Requirements in Activity-Dependent Critical Period Neural Circuit Refinement. Curr. Biol. 27, 2318–2330.e3.

30. Baltruschat, L., Prisco, L., Ranft, P., Lauritzen, J.S., Fiala, A., Bock, D.D., and Tavosanis, G. (2021). Circuit reorganization in the Drosophila mushroom body calyx accompanies memory consolidation. Cell Rep. 34, 108871.

31. Sugie, A., Marchetti, G., and Tavosanis, G. (2018). Structural aspects of plasticity in the nervous system of Drosophila. Neural Develop. 13, 14.

32. Elkahlah, N.A., Rogow, J.A., Ahmed, M., and Clowney, E.J. (2020). Presynaptic developmental plasticity allows robust sparse wiring of the Drosophila mushroom body. eLife 9, e52278.

33. Hattori, D., Aso, Y., Swartz, K.J., Rubin, G.M., Abbott, L.F., and Axel, R. (2017). Representations of Novelty and Familiarity in a Mushroom Body Compartment. Cell 169, 956–969.e17.

34. Devineni, A.V., Sun, B., Zhukovskaya, A., and Axel, R. (2019). Acetic acid activates distinct taste pathways in Drosophila to elicit opposing, state-dependent feeding responses. eLife 8, e47677.

35. Devineni, A.V., Deere, J.U., Sun, B., and Axel, R. (2020). Individual bitter-sensing neurons in Drosophila exhibit both ON and OFF responses that influence synaptic plasticity. bioRxiv, 2020.08.25.266619.

36. Guizar-Sicairos, M., Thurman, S.T., and Fienup, J.R. (2008). Efficient subpixel image registration algorithms. Opt. Lett. 33, 156.

37. Grabe, V., Baschwitz, A., Dweck, H.K.M., Lavista-Llanos, S., Hansson, B.S., and Sachse, S. (2016). Elucidating the Neuronal Architecture of Olfactory Glomeruli in the Drosophila Antennal Lobe. Cell Rep. 16, 3401–13.

38. Li, J., Ellis, K.E., and Caron, S.J.C. (2021). Photo-labeling neurons in the Drosophila brain. STAR Protoc. 2, 100381.

39. Schindelin, J., Arganda-Carreras, I., Frise, E., Kaynig, V., Longair, M., Pietzsch, T., Preibisch, S., Rueden, C., Saalfeld, S., Schmid, B., et al. (2012). Fiji: an open-source platform for biological-image analysis. Nat. Methods 9, 676–682.

40. Wan, Y., Otsuna, H., Chien, C.-B., and Charles, H. (2012). FluoRender: An Application of 2D Image Space Methods for 3D and 4D Confocal Microscopy Data Visualization in Neurobiology Research. 2012 IEEE Pac. Vis. Symp., 201–208.

41. Wan, Y., Otsuna, H., Holman, H.A., Bagley, B., Ito, M., Lewis, A.K., Colasanto, M., Kardon, G., Ito, K., and Hansen, C. (2017). FluoRender: joint freehand segmentation and visualization for many-channel fluorescence data analysis. BMC Bioinformatics 18, 280.

42. Longair, M.H., Baker, D.A., and Armstrong, J.D. (2011). Simple Neurite Tracer: open source software for reconstruction, visualization and analysis of neuronal processes. Bioinformatics 27, 2453–2454.

